# Clinical data specification and coding for cross-analyses with omics data in autoimmune disease trials

**DOI:** 10.1101/360719

**Authors:** Lorenzon Roberta, Drakos Iannis, Claire Ribet, Sophie Harris, Cordoba Maeva, Tran Olivia, Dasque Eric, Cacoub Patrice, Hartemann Agnes, Bodaghi Bahram, Saadoun David, Berenbaum Francis, Grateau Gilles, Ronco Pierre, Benveniste Olivier, Mariampillai Kuberaka, Sellam Jeremie, Seksik Philippe, Rosenzwajg Michelle, Six Adrien, Bernard Claude, Aheng Caroline, Vicaut Eric, Klatzmann David, Mariotti-Ferrandiz Encarnita

## Abstract

**Objectives:** Autoimmune and inflammatory diseases (AIDs) form a continuum of autoimmune and inflammatory diseases, yet AIDs’ nosology is based on syndromic classification. The TRANSIMMUNOM trial (NCT02466217) was designed to re-evaluate AIDs nosology through clinic-biological and multi-omics investigations of patients with one of 19 selected AIDs. To allow cross-analyses of clinic-biological data together with omics data, we needed to integrate clinical data in a harmonized database.

**Materials and Methods:** We assembled a clinical expert consortium (CEC) to select relevant clinic-biological features to be collected for all patients and a cohort management team comprising biologists, clinicians and computer scientists to design an electronic case report form (eCRF). The eCRF design and implementation has been done on OpenClinica, an open-source CFR-part **11** compliant electronic data capture system.

**Results:** The CEC selected 865 clinical and biological parameters. The CMT selected coded the items using CDISC standards into 5835 coded values organized in 28 structured eCRFs. Examples of such coding are check boxes for clinical investigation, numerical values with units, disease scores as a result of an automated calculations, and coding of possible treatment formulas, doses and dosage regimens per disease.

**Discussion:** 21 CRFs were designed using OpenClinica v3.14 capturing the 5835 coded values per patients. Technical adjustment have been implemented to allow data entry and extraction of this amount of data, rarely achieved in classical eCRFs designs.

**Conclusions:** A multidisciplinary endeavour offers complete and harmonized CRFs for AID clinical investigations that are used in TRANSIMMUNOM and will benefit translational research team.

## 1 BACKGROUND AND SIGNIFICANCE

Autoimmune and auto-inflammatory diseases (AID) are the third cause of morbidity and mortality in the world ^1^. The development of more effective and better tolerated treatments for these chronic and severely disabling diseases is an important public health issue. Recently, genetic studies have highlighted altered biological processes that are common to several AIDs^2^, and others studies have shown that an imbalance between effector T cells and regulatory T cells resulting in the rupture of immune tolerance is associated with AIDs ^3–6^. This collection of evidence is in line with the proposed reclassification of AIDs to form a continuum of diseases ranging from pure autoimmune to pure inflammatory diseases with a number of diseases displaying variable degrees of both autoimmune and inflammatory disorders ^7^. This is further sustained by immune markers common to several diseases, such as cytokines, which are currently targeted in therapeutics ^8,9^. The complexity of these diseases, due to the various genetic and environmental factors as well as patient heterogeneity, prompted the scientific community to reconsider research practices with a view to a more integrative approach. In particular, AIDs are associated with multiple and variable immune-related disorders, including dysregulation of the innate immune response or of the adaptive immune response or of both. Systems biology approaches raise the hope that a more comprehensive understanding of cells and tissues in health and disease will open up new avenues for the treatment of patients ^10,11^. These approaches will transform disease taxonomies from syndromic classification to molecular classification, and their combination, and will allow physicians to select optimal therapeutic regimens for individual patients ^12^. Recent studies have successfully identified molecular signatures associated with specific autoimmune diseases ^4,13–16^ as well as in physiological and pathological contexts ^17–19^.

Those results led us to setup an observational clinical trial, TRANSIMMUNOM, (NCT02466217) the main goal of which was to revisit the nosology of AIDs through a systems immunology approach. TRANSIMMUNOM participants include patients diagnosed with one out of 19 selected AIDs or one out of 5 control diseases (Figure 1), and healthy donors with no history of autoimmune disorders. The systems immunology approach used a multi-scale deep immunophenotyping on peripheral blood (including transcriptome, TCR repertoire, cytokine expression) and microbiome NGS studies. Importantly, classic routine biology assays as well as clinical investigations are fully part of the data collection strategy. Our aim was to integrate all these data (biology, routine biology and clinical data) so as to allow further cross-analysis of all patients and data to better characterize the immunome of each patient regardless of the initial diagnosis. A similar strategy was initiated by the National Institute of Allergy and Infectious Diseases (NIAID) under the Human Immunology Project Consortium.

**Figure 1:**
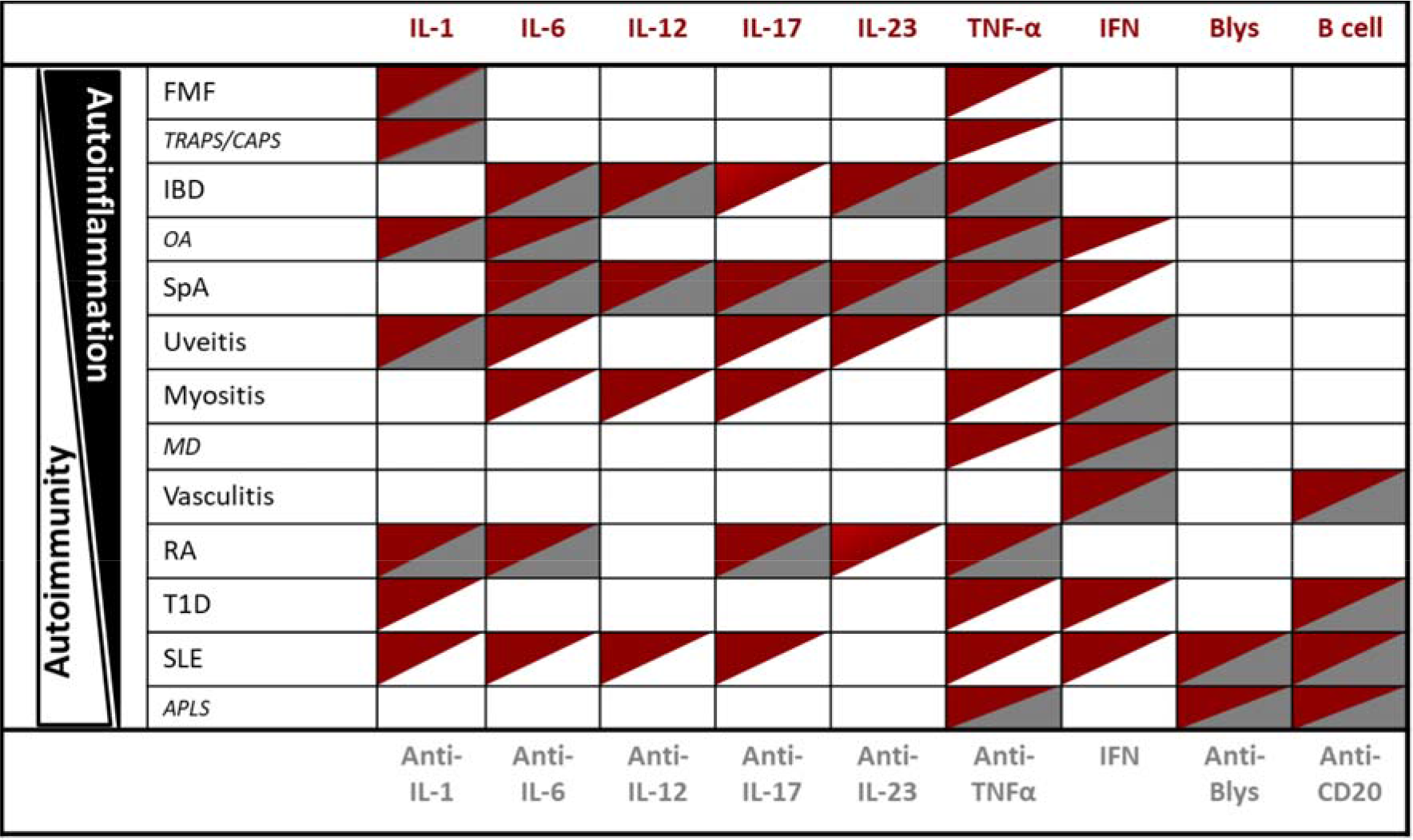
TRANSIMMUNOM selected AIDs and control diseases share immune markers and therapeutic strategies. This table shows the list of AIDs selected for the TRANSIMMUNOM trial, belonging to the AID continuum and their association with cytokines modulation (red) as well as immunotherapies targeting immune markers (grey). Abbreviation legend: Diseases: APLS anti-phospholipid syndrome, CAPS cryopyrin associated periodic syndromes, FMF familial mediterranean fever, IBD - inflammatory bowel disease, MD muscular dystrophy, OA osteoarthritis, RA rheumatoid arthritis, SLE systemic erythematosus lupus, SpA spondyloarthritis, T1D type 1 diabetes, TRAPS tumor necrosis factor receptor-associated periodic syndrome. Cytokines: IFN interferon, IL-1 interleukin-1, IL-6 interleukin-6, IL-12 interleukin-12, IL-17 interleukin-17, IL-23 interleukin-23, TNF-α tumor necrosis factor alpha. Immunotherapies: Anti–BLyS anti-BLyS monoclonal antibody, Anti–CD20 anti-CD20 monoclonal antibody Anti–IL-1 anti-interleukin-1 monoclonal antibody, Anti–IL-6 anti -interleukin-6 monoclonal antibody, Anti–IL-12 anti-interleukin-12 monoclonal antibody, Anti-IL-17 anti-interleukin-17 monoclonal antibody, Anti–IL-23 anti-interleukin-23 monoclonal antibody, Anti-TNFα tumor necrosis factor alpha-blockers.

## 2 OBJECTIVES AND OUR CONTRIBUTION

Therefore, we needed to develop data integration approaches to efficiently record and store collected data such that we could easily analyze them afterwards through computational biology approaches. The first challenges of the project were to implement a comprehensive case report form (CRF) covering all diseases in terms of clinical data and biomarkers and to provide a user-friendly, vocabulary-controlled and not expensive platform with standard vocabulary to record all data collected by the clinical assistant during patient interviews. To meet these challenges, we assembled the multidisciplinary “Cohort Management Team (CMT)” composed of clinicians from different specialties, nurses, biology medical doctors, clinical trial methodologists, immunologists and computer scientists.

Here we present our electronic CRF (eCRF), designed using an open-source electronic data capture (EDC) tool, capturing more than 5000 multiparametric coded values from 865 harmonized clinical and biological parameters per subject included in a multi-disease clinical trial focusing on 24 diseases, 22 areas of clinical investigation and one vast set of routine biology assays. Altogether, we believe that this effort could be of interest for small cohort studies for which the commercially available eCRF services are not accessible.

## 3 MATERIAL AND METHODS

### 3.1 Study population

Patients with one of the following AIDs, of the AID continuum are recruited for TRANSIMMUNOM trial (Figure 1): familial mediterranean fever (FMF), ulcerative colitis, Crohn’s disease, spondyloarthritis, uveitis, myositis (polymyositis, dermatomyositis, inclusion-body myositis, necrotizing and anti-synthetase related myositis), ANCA-related vasculatis (Churg-Strauss’ disease and granulomatosis with polyangiitis (ex Wegener), non-ANCA-related vasculitis (such as Behçet’s disease, cryoglobulinaemia and Takayasu), rheumatoid arthritis (RA), type-1-diabetes and systemic lupus erythematosus (SLE). We also included patients with diseases exhibiting symptoms similar to those of some AIDs but linked to different gene mutations (control diseases), such as TRAPS and CAPS as a control for FMF, or diseases with a similar autoimmune mechanism with overlapping clinical/biological features, such as antiphospholipid syndrome (APLS) as a control of for SLE, or degenerative diseases that do not have the same mechanism as AIDs such as osteoarthritis for RA and muscular dystrophy for myositis. Finally, healthy volunteers are included.

### 3.2 Cohort Management Team

Set up to interact with a Clinical Expert Consortium (CEC), a Cohort Management Team (CMT) of biological experts, routine laboratory personnel, clinical trial methodologists and clinical investigation centre harmonized the clinical and laboratory outcomes/results. The CMT ensured that all required data are collected in an appropriate format for analyses and that the questions are unambiguous. The computer scientist defined the data and metadata structure required to minimize non-controlled data entry and to specify the expected values. The overall design was supervised by an immunologist involved in the scientific part of the clinical trial, who liaised between the clinicians and the computer scientist.

### 3.3 Data collection for eCRF design and coding

Each clinician received an Excel form to be filled in with the description of the item to be recorded in a standardized manner: item ID, item value type (string, decimal); list of predetermined item values; item value unit (if applicable); item value range (if applicable). Afterwards, all the data collected from the different specialties were grouped and harmonized using CDISC standards.

### 3.4 OpenClinica implementation

Given the amount of data to be collected across 19 AIDs and 5 control diseases, OpenClinica v3.14, an open-source CFR-part 11 compliant electronic data capture platform has been selected for the design and capture of selected clinico-biological data. A test and production instances have been installed on dedicated and secured CentOS virtual machines with 16Go RAM, 8 cores and 15 Go disk space each. OpenClinica’s application server (Tomcat v9.0.6) and database server (PostgreSQL v.9.5) parameters have been upgraded to fit multiple simultaneous data entry and data extraction, in particular JAVA_OPTS for heap memory have been upgraded to 8 Go instead of 1Go Mo by default.

### 3.5 Patient anonymization

To have completely anonymized subjects who are also unique (no double entries for the same subject in our database because of anonymity) we developed the Anonymized Subject Unification (ASU) system as a completely autonomous system that can be used for any clinical trial. Briefly, ASU takes advantage of a unique identifier of each subject (like Paris Hospital patient number [NIP] or French healthcare registration number [INSEE]) to produce a simple 4-letter code by using a one-way encryption technique.

## 4 RESULTS

### 4.1 OpenClinica as the compromise in designing a multi-disease eCRF

The TRANSIMMUNOM observational trial targeted recruitment of 1,000 patients suffering from one out of 24 diseases and healthy controls. During a single visit, patient medical history and clinical investigations are performed together with the collection of samples (blood, serum and feces) for further multi-omics analyses. The goal of the trial is to revisit the nosology of AIDs by defining groups/clusters of patients based on clinical and molecular signatures that cut across disease classification. To deal with the expected amount of heterogeneous (such as disease severity scores, imaging data, biological measures) from routine clinical investigations, and to allow the cross-evaluation of clinical and omics data, we needed to develop an eCRF with a system that allows further omics data integration. We selected OpenClinica (OC) as an electronic data capture (EDC) tool to support our eCRF design. OC is an open-source CRF-part 11 compliant EDC able to design complex eCRFs for large studies ^20,21^. One of the major features of OC is to rely on Clinical Data Acquisition Standards Harmonization (CDASH) from the Clinical Data Interchange Standards Consortium (CDISC) ^22^, which allows the harmonization of clinical and biological data coding. Finally, OC includes the mandatory validation of all recorded data to ensure data quality ^20^. In addition, the main strategy of TRANSIMMUNOM is to cross-analyze data from multiple AIDs, each of which is usually characterized by particular clinical investigation records and biological data measures. We anticipated the final cross-analysis, which would require the same information for all the diseases. Finally, the eCRF had to follow regulatory guidelines and Good Clinical Practices to ensure data entry, traceability and integrity throughout the patient recruitment period. Although installation and implementation of OC is not trivial, as it requires computer science expertise and time, we decided to favour the landscape of possibilities offered by OC to fulfil our study requirement.

### 4.2 A multidisciplinary workflow ensuring the design of a robust multi-disease eCRF

Expertise in different but converging fields was pooled in the CMTs, each of which participated in a 3-step workflow to (1) define the protocol, (2) design and (3) validate the eCRF (Figure 2). The first step of protocol definition involved a Clinical Expert Consortium (CEC) to define the list of items for all the patients with the aim of collecting exactly the same information regardless of the disease. All the clinical specialists together selected a sample of items per specialty so that the CRF was reasonably comprehensive and synthetic. Biology lab experts were also questioned to ensure the feasibility of sample drawing and of the required biology assays. Upon collection and validation of the actual items to be recorded, the specification of the database started with the design of an e-template where the computer scientist structured the information for each item by imposing the format of the data and metadata. Once the e-template was defined, we proceeded to the eCRF design: the CEC, in close collaboration with the computer scientist, designed the clinical coding of clinical investigation data following an unambiguous format for each item with maximized use of a predefined list of responses in order to avoid erroneous data entry. Biology lab experts defined for each parameter measured the value type (string, integer, decimal, Boolean), as well as the unit and range, when applicable. All the information was summarized in a spreadsheet and converted by the computer scientist into a PostgreSQL relational database following the OpenClinica structure. Finally, clinical research technicians evaluated the user-friendliness of the eCRF, the clinical research assistant evaluated the item relationship constraints, and finally the CMT validated the eCRF with a patient “Zero” simulation before release for production.

**Figure 2:**
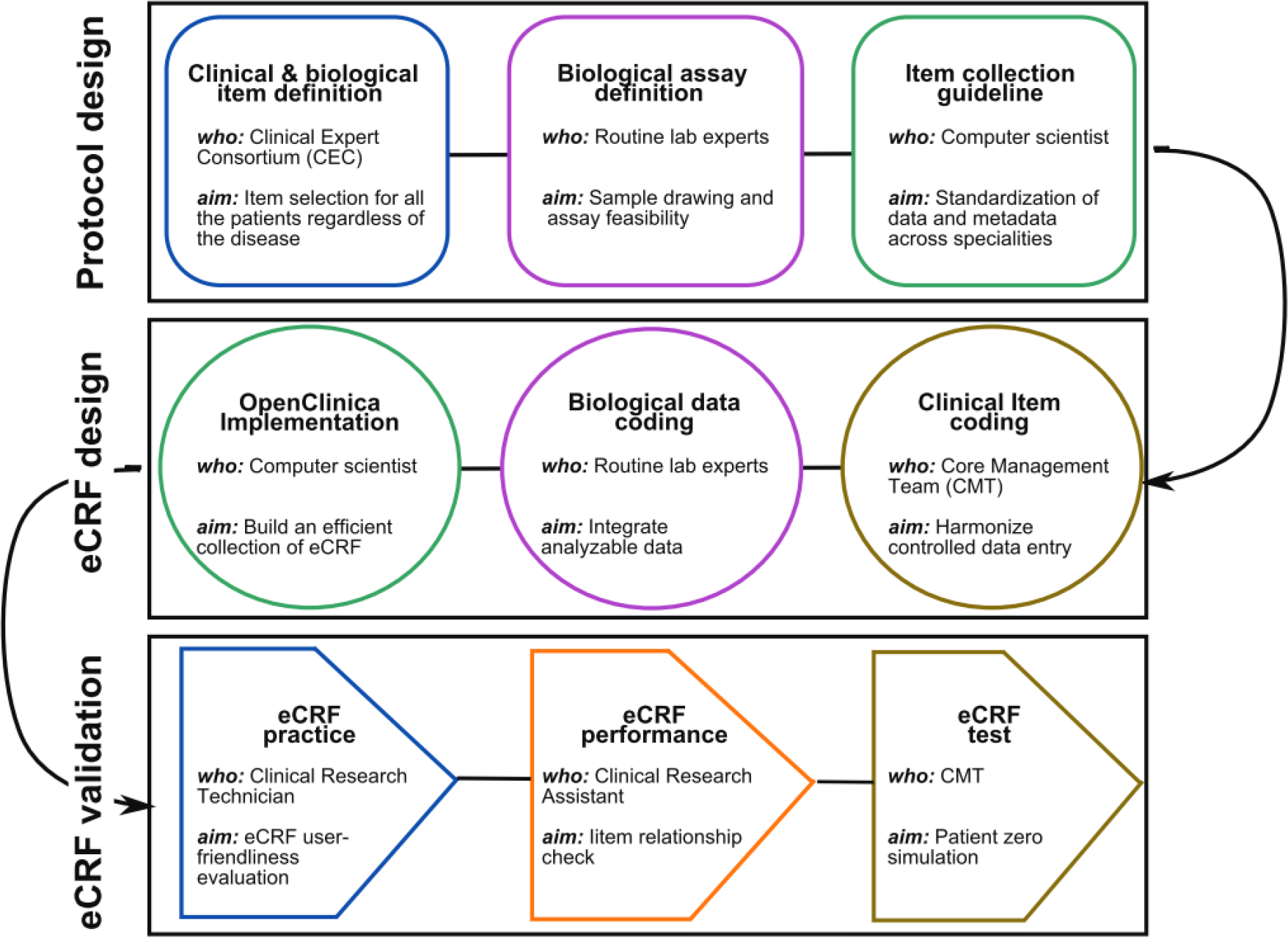
eCRf workflow. The figure represents the 3-steps workflow adopted for the eCRF design and implementation: (1) Protocol design, (2) eCRF design and (3) eCRF validation. In each box are listed the actions, its aim and the person in charge of it. Color code: Blue: clinical team, purple: biological team, green: computer scientist, and orange: the trial monitor team and brown the Core management team (see methods for description).

### 4.3 An integrated multi-disease eCRF

As AIDs belong to different medical specialities, the CEC comprised clinicians working in rheumatology, internal medicine, gastroenterology, diabetology, ophthalmology, medical biology, nephrology and genetics who ensured the feasibility of data collection in terms of cost, patient morbidity and examination invasiveness. The list of information to be collected for all the participants was organized in 4 categories. For each recorded item, we defined the type of value such as free text field, free numerical field, automated calculation, check-box, drop-down list and calendar/date field (Figure 3). The first group of CRFs was built under the “Patient description” category and included classic clinical information required to assess the biology and social environment of the patient. Altogether, we selected 70 items organized as 7 CRFs (Figure 3A & Supplementary material). Each CRF collects 4 to 30 different items. The second set of CRFs focuses on “Common clinical monitoring” and was organized as 5 CRFs collecting generic clinical data at the day of the visit and accounting in all for 88 items (Figure 3B & Supplementary material). The third category explore the “Routine biology monitoring” (Figure 3C, Supplementary material & Table 1) and covered a wide spectrum of tests.

**Figure 3:**
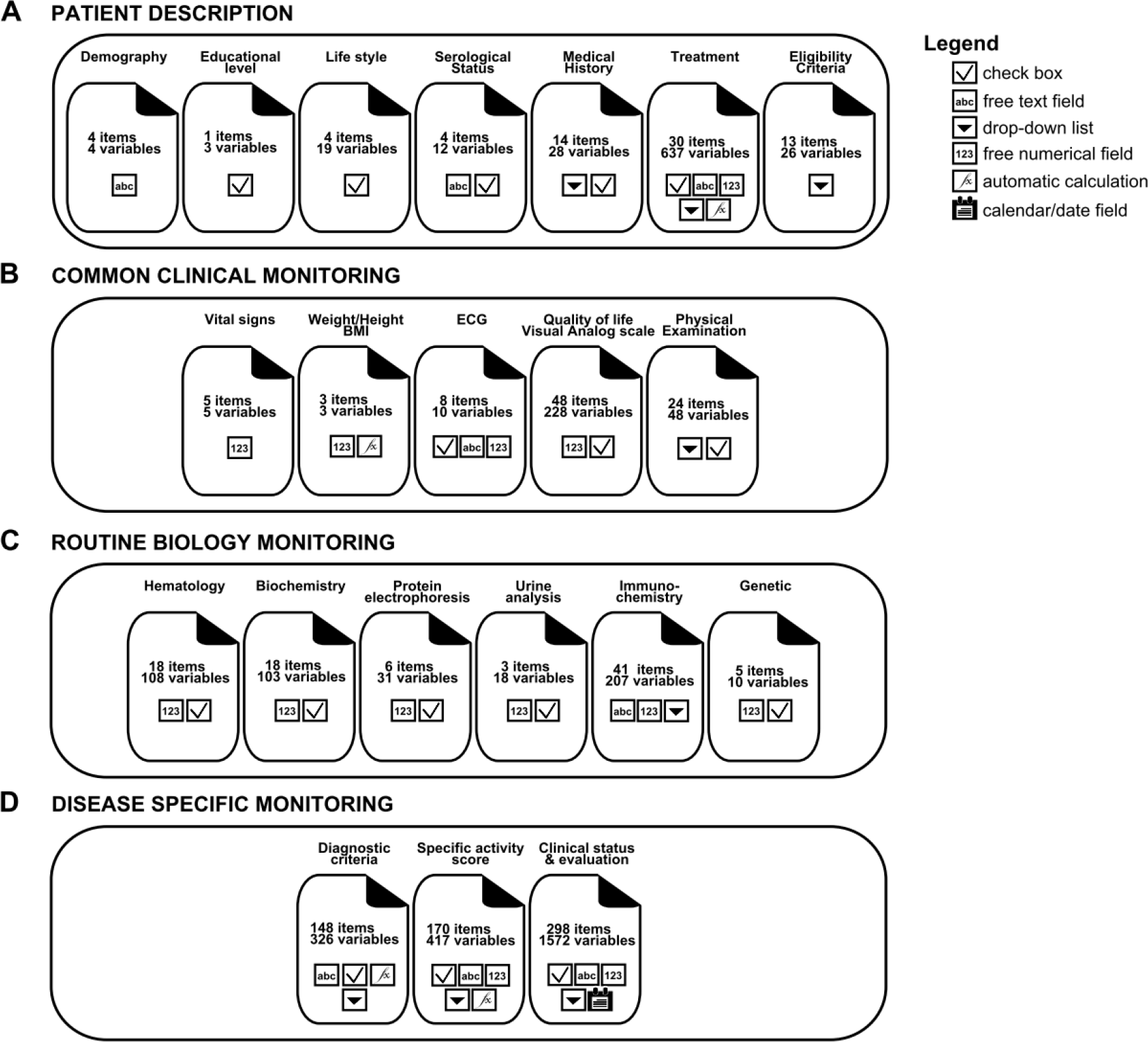
Schematic representation of the TRANSIMMUNOM integrated eCRF. Four categories of eCRFs were designed (A-D). Each category is composed of several eCRFs (form icon), each of which contained the indicated number of items for which 1 to 8 variables were coded. The type of values are indicated in the square boxes (see legend), so as to check-box, free text field, drop-down list, free numerical field, automatic calculation and calendar/date field. Altogether, 865 items were coded resulting in 5835 variables organized in 21 eCRFs. See Supplementary material for details on eCRF

**Table 1:**
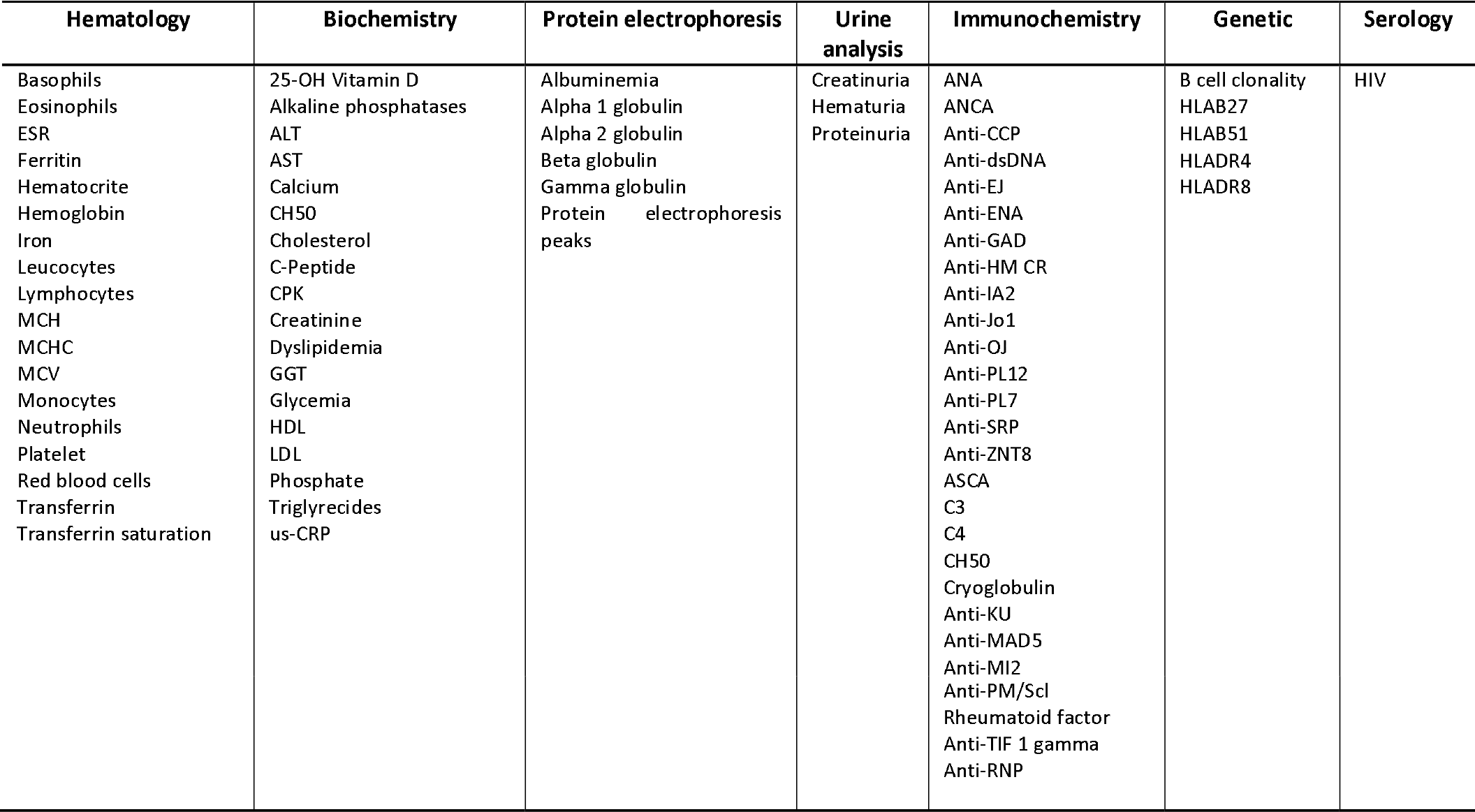
List of routine biology assay in the TRANSIMMUNOM trial. Table abbreviation legend: ALT - alanine aminotransferase, ANA - antinuclear antibodies, ANCA - anti-neutrophil cytoplasmic antibodies, Anti-CCP - anti-cyclic citrullinated peptide antibodies, Anti-dsDNA - Anti-double stranded DNA antibodies, Anti-EJ - anti-glycyl-transfer RNA synthetase antibodies, Anti-ENA - anti-extractable nuclear antigens antibodies, Anti-GAD - anti-glutamic acid decarboxylase antibodies, Anti-HMGCR -anti-3-hydroxy-3-methylglutaryl-coenzyme A reductase antibodies, Anti-IA2 - anti-Islet antigen-2 antibodies, Anti-Jo1 - anti-histidyl tRNA synthetase antibodies, Anti-Ku - anti- Ku antigen antibodies, Anti-MDA5 -melanoma differentiation-associated gene 5 antibodies, Anti-MI2 - anti–Mi-2 antibodies, Anti-OJ - anti-isoleucyl-tRNA synthetase antibodies, Anti-PL7 - anti-threonyl-tRNA synthetase antibodies, Anti-PL12 - anti-alanyl-tRNA synthetase antibodies, Anti-PM/Scl - anti- nucleolar macromolecular complex PM/Scl, Anti-RNP -anti-nuclear ribonucleoprotein antibodies, Anti-SRP - anti-signal recognition particle antibodies, Anti-TIF1-gamma – anti-transcriptional intermediary factor 1-gamma antibodies, Anti-ZnT8 -anti-zinc transporter 8 antibodies, ASCA - anti-Saccharomyces cerevisiae antibodies, AST - aspartate aminotransferase, C3 -complement fraction 3, C4 - complement fraction 4, CH50 - total complement activity, CPK - Creatinine phosphokinase, ESR - erythrocyte sedimentation rate, GGT - gamma-glutamyl transferase, HDL - high-density lipoprotein, HIV - Human Immunodeficiency Virus, HLA-B27-B51-DR4-DR8 Human leukocyte antigen -B27-B51-DR4-DR8, LDL - low-density lipoprotein, MCH - mean corpuscular haemoglobin, MCHC - mean corpuscular hemoglobin concentration, MCV - mean corpuscular volume, us-CRP - ultrasensitive c-reactive protein.

These included biological assessment of organ function (liver, kidney, bone marrow) and of inflammation state and safety, organized in 6 CRF and covering 91 parameters. Finally, the last set of CRFs recorded “Disease-specific monitoring” data and was subdivided into 3 CRFs (Figure 3D & Supplementary material) capturing 616 items, including disease activity scores as described in Table 2. This is thought to be as wide as possible in identifying clinical parameters not usually collected in a particular disease including imaging and histology features to allow the identification of disease profile, disease severity and features possibly shared by diseases. Each clinician of the CEC identified a collection of features observed in his/her specialty as classic or rare parameters. The CMT gathered all the parameters from the different specialties and listed them in the clinical status and clinical evaluation CRFs. Altogether, we selected 865 parameters to describe each patient regardless of the disease.

**Table 2:** Disease specific diagnostic criteria and activity score.

| Disease | Diagnostic criteria | Activity score |
| --- | --- | --- |
| Familial Mediterranean Fever/TRAPS/CAPS | Heller<sup>23</sup><br>*Gene mutation: MEFV; TNFRSF1; NLRP3* | AIDAI<sup>24</sup> |
| Ulcerative colitis | clinical and histological features | Mayo<sup>25</sup> |
| Crohn's disease | clinical and histological features | HBI<sup>26</sup> |
| Spondyloarthritis | ASAS <sup>27</sup><br>Modified New-York criteria <sup>28</sup> | BASDAI <sup>29</sup> |
| Uveitis | non-infectious uveitis | NA |
| Myositis/dystrophy | clinical and biological | NA |
| Vasculitis | <u>Behcet's disease</u><br>ICBD <sup>30</sup><br><u>Churg-Strauss</u><br>ACR <sup>31</sup><br><u>Cryoglobulinemia</u> <sup>32</sup><br><u>Wegener</u><br>ACR <sup>33</sup><br><u>Takayasu</u><br>ACR <sup>34</sup> | BVAS <sup>35</sup><br>NIH <sup>36</sup> |
| Rheumatoid Arthritis | ACR; EULAR <sup>37</sup> | DAS-28 <sup>39</sup> |
| Osteoarthritis | Kellgren-Lawrence <sup>38</sup> |  |
| Type-1-Diabetes | ADA <sup>40</sup> | IDAA1C <sup>41</sup> |
| Systemic Lupus Erythematosus / | ACR <sup>42,43</sup> | SLEDAI <sup>45,46</sup> |
| Anti-phospholipid syndrome | Sapporo <sup>44</sup> |  |

### 4.4 Clinical coding and CDASH harmonization

Because of the heterogeneity of the selected parameters, clinical coding was designed as an unambiguous format based on CDASH standards with maximized use of a predefined list of responses, and was developed as a pragmatic, clinically-validated medical terminology with an emphasis on ease-of-use data entry, retrieval and data analysis. We therefore defined and validated for each parameter, wherever possible, the data-type (numerical, text, date, predefined lists of options, value ranges) and units (when applicable) for all the parameters identified in order to harmonize the information regardless of the collection time and person and to avoid errors due to mistyping. Examples are Yes/No check boxes for clinical investigation, numerical values with a list of relevant units according to the parameter, disease scores as a result of the automated sum of several scores, treatment description including the coding of possible formulas, doses and dosage regimens (Table 3). We then coded all the possible/expected values that each item could take and identified 1 to 8 possible variables per item coded as one of the value type. This work was especially critical for the description of patient treatments. The list of all possible treatments regimen within each specialty was fully generated with clinicians and is available in the database as a menu list of 637 variables. Altogether, we built a database with 3815 uniquely coded variables. However, since clinical status and evaluation of several diseases share identical CRFs, we reached 5835 possible variables per patient. Altogether, we designed a collection of 21 CDASH harmonized CRFs recording 865 parameters with 5835 coded variables systemically for all the patients and healthy donors included in the TRANSIMMUNOM trial.

**Table 3:**
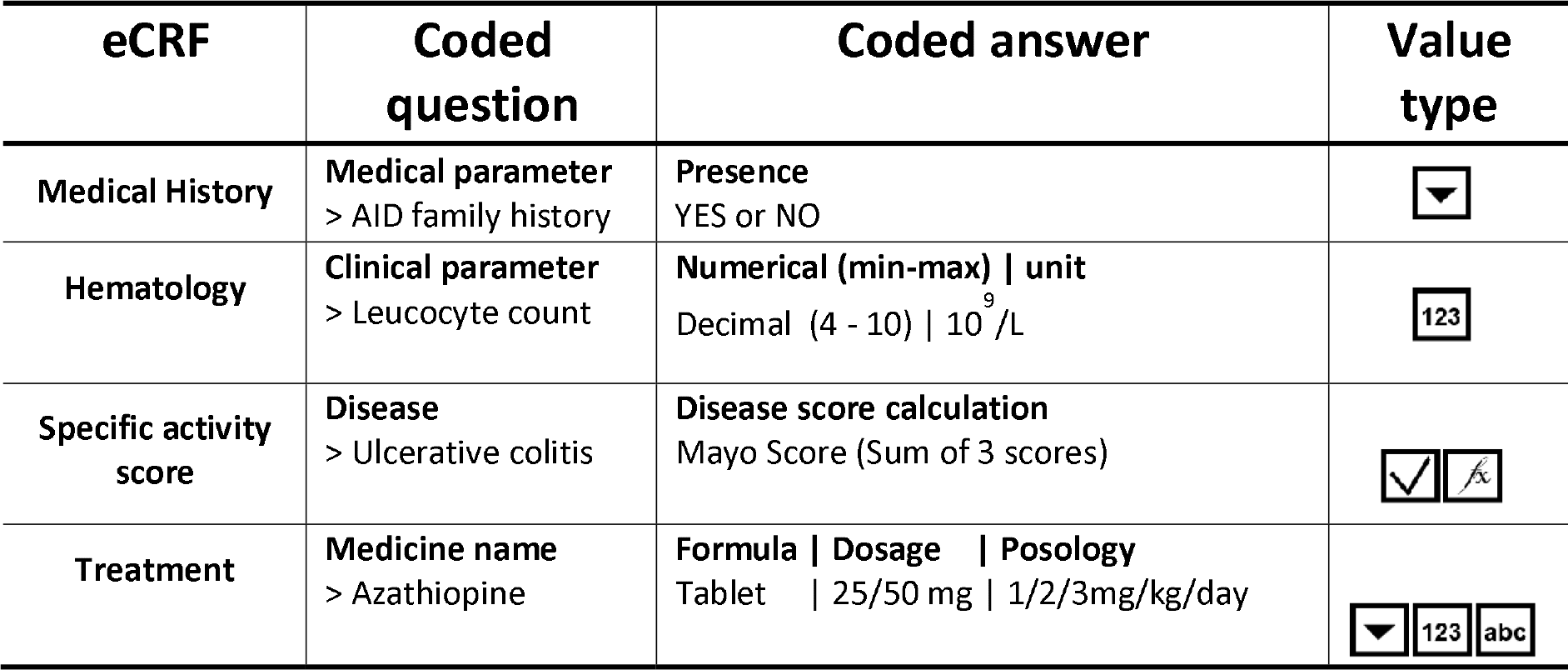
Clinical item coding. For 5 exemplary eCRF, a coded question and expected coded answer are described with the type of values to be entered following Figure 3 legend.

## 5 DISCUSSION

Clinical data management is of utmost importance for any clinical study. This includes clinical information collection, validation and storage, usually completed through the use of CRFs. While generally clinical research organizations (CROs) propose and use eCRFs, most academic sponsored clinical studies still take advantage of the cost-benefit of paper versions of CRFs, sometimes in combination with Excel based databases^20^. However, such tools, although convenient, lack validation and data traceability. In addition, they do not usually use harmonized vocabulary and allow free text data entry. These drawbacks were particularly counterproductive in our multi-disease clinical trial from several points of view. First, the main goal of our trial is cross-analysis of multi-omics data obtained with clinical and lab biology data from 1,000 patients selected for one out of 24 AIDs or control diseases. Therefore, we needed an efficient and homogenized set of clinical and routine biology lab for all the patients, which led to the selection of 865 parameters and to the coding of more than 5000 values. This vast amount of data would have been unmanageable using classic paper CRFs and spreadsheets. Second, the amount of information to be collected requires a thorough validation with automated rules and limited free text data entry to avoid mistyping and errors. Again, this cannot be handled using classic methods. Third, for cross-analysis, we need to be able to extract clinical and routine biology data efficiently so that we can filter for parameters as variables of interest (such as gender, BMI, disease activity, autoantibody level). Again, considering the number of patients to be recruited in the TRANSIMMUNOM trial, this would have been impossible. And finally, as regards the disease heterogeneity, it would have been too expensive and complicated to ask an eCRF provider to design such integrated CRFs. For all these reasons, we decided to take advantage of an open-source EDC, Open-Clinica, for the implementation of our eCRFs. Although we anticipated that the design and computer-based requirements would be time-consuming, we found in this tool a number of advantages that allow (i) the integration of a very significant amount of multi-parametric data, (ii) the possibility to design constraints rules between entries to control data entry errors, associated with red flags in the case of errors (for instance a man cannot be pregnant), (iii) the validation of the data entry by a third person who double-checks (the latter advantages being CFR 21 - part 11 compliant) and (iv) the addition of short instructions on the CRF page when needed to guide the data entry and explain to the investigator how to fill in the eCRFs.

Altogether, this choice allowed the design of a controlled series of CRFs using harmonized vocabulary to record data across 19 AID patients, 5 control disease patients and healthy donors. This was made possible by the workflow we dedicated to the project, going from the selection of parameters to be collected for all patients regardless of the disease to the coding of all possible values per parameter in a harmonized manner based on CDASH coding. 26 persons were involved in the process, including 14 clinicians, 1 computer scientist, 7 scientists, 3 clinical research technician and assistant as well as 2 medical biologists for more than 100 hours of meetings and discussion over a year and a half. Clinical data coding has the enormous advantages that it (i) pools reported terms in medically meaningful groups, (ii) facilitates identification of common data sets for evaluation of clinical information, (iii) supports consistent retrieval of specific cases or medical conditions from a clinical database and (iv) smooths electronic data interchange of clinical safety information.

Finally, our CRFs covers a wide spectrum of clinical and routine biology data of interest for most AIDs, offering the community a pre-designed set of CRFs that can be used together or individually. Although clinical safety was not added to our set of CRFs, because of the noninterventional nature of the TRANSIMMUNOM trial, this could easily be done if needed. This complex set of data has been harmonized and the database designed to store and query efficiently the massive amount of data stored. Altogether, a truly multidisciplinary endeavour led to the design and implementation a collection of 21 CRFs capturing more than 5000 coded values that are now used in TRANSIMMUNOM and could benefit the academic clinical community studying AIDs.

## Acknowledgments

We are grateful to Frédéric Mariotti as an informatic subcontractor for helping in the implementation and maintenance of the OpenClinica instance and to the OpenClinica community, in particular Gerben Rienk Visser from Trial Data Solution for providing help in OpenClinica parameter setup.

## Funding

The work of RL, ID, CR, SH, CA, DK, EMF is funded by the LabEx Transimmunom (ANR-11-IDEX-0004-02) as well as by Assistance Publique-Hôpitaux de Paris and Sorbonne Université. TRiPoD funded the TCR-relevant part of the study.

## Author contributions

RL, ID, CR, CA, AS, DK and EMF composed the Cohort Management Team (CMT) for the design and implementation of the eCRF. RL, ED, PC, AH, BB, DS, FB, GG, PR, OB, KM, JS, PS, MR, CB formed the Clinical Expert Consortium (CEC) and defined the selected clinical and biological data. RL and CR wrote the CRF in agreement with the CMT recommendations. ID performed the computer science part of the work. CM, SH, TO worked on the eCRF validation. EMF coordinated the design and implementation. EMF and RL wrote the manuscript with input from all authors. EMF and DK conceived and supervised the entire work.

## Competing interests

Authors have no competing interests to declare.

